# Mechanical loading reveals an intrinsic cardiomyocyte stiffness contribution to diastolic dysfunction in murine cardiometabolic disease

**DOI:** 10.1101/2024.02.21.581448

**Authors:** Johannes V. Janssens, Antonia J.A. Raaijmakers, Parisa Koutsifeli, Kate L. Weeks, James R. Bell, Jennifer E. Van Eyk, Claire L. Curl, Kimberley M. Mellor, Lea M.D. Delbridge

## Abstract

Cardiometabolic syndromes including diabetes and obesity are associated with occurrence of heart failure with diastolic dysfunction. There are no specific treatments for diastolic dysfunction, and therapies to manage symptoms have limited efficacy. Understanding of the cardiomyocyte origins of diastolic dysfunction is an important priority to identify new therapeutics. The investigative goal was to experimentally define *in vitro* stiffness properties of isolated cardiomyocytes derived from rodent hearts exhibiting diastolic dysfunction *in vivo* in response to dietary induction of cardiometabolic disease.

Male mice fed a high fat/sugar diet (HFSD vs control) exhibited diastolic dysfunction (echo E/e’ doppler ratio). Intact paced cardiomyocytes were functionally investigated in three conditions: non-loaded, loaded and stretched.

Mean stiffness of HFSD cardiomyocytes was 70% higher than control. E/e’ for the origin hearts was elevated by 35%. A significant relationship was identified between *in vitro* cardiomyocyte stiffness and *in vivo* dysfunction severity. With conversion from non-loaded to loaded condition, the decrement in maximal sarcomere lengthening rate was more accentuated in HFSD cardiomyocytes (vs control). With stretch, the Ca^2+^ transient decay time course was prolonged. With increased pacing, cardiomyocyte stiffness was elevated, yet diastolic Ca^2+^ elevation was attenuated. Our findings show unequivocally that cardiomyocyte mechanical dysfunction cannot be detected by analysis of non-loaded shortening.

Collectively, these findings demonstrate that a component of cardiac diastolic dysfunction in cardiometabolic disease is derived from cardiomyocyte stiffness. Differential responses to load, stretch and pacing suggest that a previously undescribed alteration in myofilament-Ca^2+^ interaction contributes to intrinsic cardiomyocyte stiffness in cardiometabolic disease.

**KEY POINTS:**

- Understanding cardiomyocyte stiffness components is an important priority for identifying new therapeutics for diastolic dysfunction, a key feature of cardiometabolic disease.
- In this study cardiac function was measured *in vivo* (echocardiography) for mice fed a high-fat/sugar diet (HFSD, ≥25weeks). Performance of intact isolated cardiomyocytes derived from the same hearts was measured during pacing under non-loaded, loaded and stretched conditions *in vitro*.
- Calibrated cardiomyocyte stretches demonstrated that stiffness (stress/strain) was elevated in HFSD cardiomyocytes *in vitro* and correlated with diastolic dysfunction (E/e’) *in vivo*. HFSD cardiomyocyte Ca^2+^ transient decay was prolonged in response to stretch. Stiffness was accentuated with pacing increase while the elevation in diastolic Ca^2+^ was attenuated.
- Data show unequivocally that cardiomyocyte mechanical dysfunction cannot be detected by analysis of non-loaded shortening.
- These findings suggest that stretch-dependent augmentation of the myofilament-Ca^2+^ response during diastole partially underlies elevated cardiomyocyte stiffness and diastolic dysfunction of hearts of animals with cardiometabolic disease.

Abstract Figure Legend
Understanding cardiomyocyte stiffness components is an important priority for identifying new therapeutics for diastolic dysfunction, a key feature of cardiometabolic disease. In this study cardiac function was measured *in vivo* (echocardiography) for mice fed a high-fat/sugar diet (HFSD, ≥25weeks). Performance of intact isolated cardiomyocytes derived from the same hearts was measured during pacing under non-loaded, loaded and stretched conditions *in vitro.* Calibrated cardiomyocyte stretches demonstrated that stiffness was elevated in HFSD cardiomyocytes *in vitro* and correlated with diastolic dysfunction (E/e’) *in vivo*. These findings show that stiff hearts are characterized by stiff cardiomyocytes in metabolic disease.

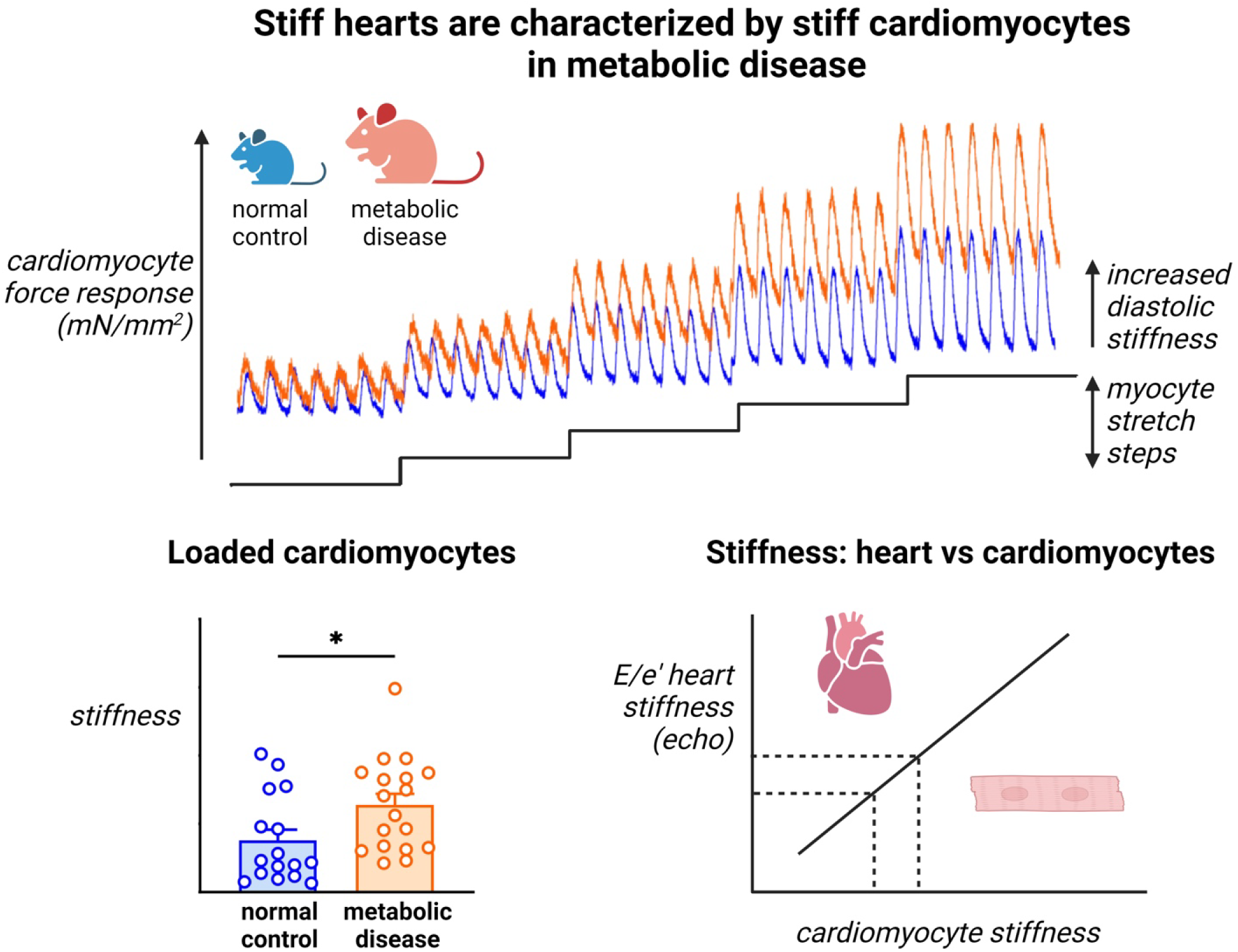

## INTRODUCTION

Increasingly, cardiometabolic syndromes, including diabetes and obesity, are associated with occurrence of heart failure with a diastolic dysfunction patho-phenotype (Sotomi *et al*., 2021). Diastolic dysfunction is characteristic of heart failure with preserved ejection fraction (HFpEF) which has now overtaken heart failure with reduced ejection fraction (HFrEF) comprising the majority of new heart failure diagnoses (McHugh *et al*., 2019; Jackson *et al*., 2022). Diastolic dysfunction is also linked with progression of diabetic cardiomyopathy (Patil *et al*., 2011; Redfield, 2016). With both HFpEF and diabetic heart disease, early development of diastolic dysfunction can be detected even when other phenotypic features/comorbidities are heterogeneous (i.e. hypertrophy, dilation, natriuretic factor elevation, hypertension) (Kittleson *et al*., 2023). Common clinical features of cardiometabolic syndromes, including obesity and insulin resistance, are independent risk factors for diastolic dysfunction as a prelude to heart failure (Rozenbaum *et al*., 2019; Kamon *et al*., 2020). The experimental induction of these phenotypes using a high/fat sugar diet feeding protocol has been shown by us and others to reliably produce diastolic dysfunction in rodents (Pulinilkunnil *et al*., 2014; Dia *et al*., 2020; Sowers *et al*., 2020; Wingard *et al*., 2021; Daniels *et al*., 2022). Cardiometabolic syndromes can involve the ‘silent’ presence and progression of diastolic dysfunction for an extended period. Symptoms can be subclinical and/or detectable only in challenge conditions (i.e. as reduced diastolic reserve in exercise) or when exacerbated by co-morbidity (Leung *et al*., 2015; Lam *et al*., 2019). Diagnosis of heart failure with diastolic dysfunction is independently associated with higher incidence of adverse clinical events (Sotomi *et al*., 2021). Notably, there is currently no specific treatment for diastolic dysfunction and/or HFpEF, and therapies to manage symptoms have limited efficacy (Delbridge *et al*., 2022).

In diastolic dysfunction, the relaxation and filling phases of the cardiac cycle are impaired. The condition is generally described qualitatively as an increase in left ventricular wall ‘stiffness’ (or by a decrease in compliance, the inverse of stiffness) (Daniels *et al*., 2022; Janssens *et al*., 2024). Clinically and experimentally, reduced compliance can be measured invasively as an elevation in left ventricular filling pressure or an increase in the gradient of the diastolic pressure-volume relationship. Non-invasively, on doppler echocardiogram, an increase in the E/e’ ratio [early (E) mitral valve blood flow rate to early mitral annulus tissue motion (e’)] is an index used to identify diastolic abnormality (Lo & Thomas, 2010).

Myocardial stiffness is commonly considered to reflect myocardial extracellular matrix abnormalities linked with increased tissue fibrosis. While a component of reduced compliance likely reflects fibrotic infiltration, diastolic dysfunction can be identified in settings where there is no evidence of increased interstitial fibrosis burden (Curl *et al*., 2018). There is growing appreciation that alterations of myocardial compliance in diastolic dysfunction disease states represents underlying intrinsic cardiomyocyte stiffness pathology which contributes to impaired relaxation and filling (Zhou & Pu, 2016; Villalobos Lizardi *et al*., 2022). Understanding the cardiomyocyte origin of diastolic dysfunction is an important priority to identify new molecular targets and to support the development of new diagnostics and therapeutics to reduce morbidity and mortality in cardiometabolic disease states.

Cardiomyocyte origins of diastolic dysfunction potentially comprise structural and operational components. Evidence of molecular changes in sarcomeric and cytoskeletal elements is extensive (Villalobos Lizardi *et al*., 2022; Loescher *et al*., 2023; Janssens *et al*., 2024). Operationally, the cardiac cycle of contraction and relaxation is driven by the cyclic flux of cardiomyocyte activator Ca^2+^ to initiate contraction (myocyte sarcomere shortening) and to allow relaxation (myocyte sarcomere lengthening). The relationship between cytosolic Ca^2+^ levels and myofilament activation is dynamic and depends on several external factors some of which include stretch and pacing (Fabiato & Fabiato, 1978; Varian & Janssen, 2007).

Conventionally, the steady-state force-Ca^2+^ analysis (typically performed in permeabilised, quiescent preparations) is used to summarise the myofilament-Ca^2+^ response and is the product of many complex myofilament protein interactions. Less work has been done to investigate the dynamic changes in the myofilament-Ca^2+^ relationship in paced intact cardiomyocytes, especially in disease contexts. In diastolic dysfunction, specifically in HFpEF animal models, evidence is accruing that Ca^2+^ deficit and impaired non-loaded shortening *are not* phenotypic (Kilfoil *et al*., 2020). Indeed cardiomyocyte myofilament hyper-activation and elevated Ca^2+^ levels have been reported (Curl *et al*., 2018; Louch, 2020; Saad *et al*., 2023). As we have recently proposed, the role for elevated diastolic Ca^2+^ levels and augmented myofilament-Ca^2+^ response comprising a cross-bridge derived component of cardiomyocyte diastolic stiffness is increasingly plausible (Janssens *et al*., 2024). This experimental enquiry focusses on intact cardiomyocyte mechanical and Ca^2+^ performance in response to load, stretch and modulated pacing conditions.

While the term ‘stiffness’ is used as a general descriptor of impaired compliance in a cardiovascular setting, it is more formally specified quantitatively in physical science disciplines (Mirsky, 1976). Stiffness is defined as the extent to which an object resists deformation. Stiffness is influenced by the geometry and the intrinsic material properties of an object. In the cardiac literature and in this study, stress/strain relations are calculated to assess cardiomyocyte resistance to stretch. Stress is defined as force per cross-sectional area, and strain is the application of a normalized stretch or deformation. The ratio of stress/strain defines the cardiomyocyte resistance to stretch independent of geometry and is known as ‘Young’s Modulus’. While stiffness and Young’s Modulus are not equivalent, the term ‘stiffness’ is commonly used (even though Young’s Modulus is the correct formal term). In the cardiac setting, a stretch or deformation state is commonly described as ‘loading’ – for example ‘pre-load’ (diastolic stretch) and ‘after-load’ (resistance to shortening). Depending on the experimental design, stiffness behaviour of myocardial tissues, active cardiomyocytes and/or myocyte sub cellular cytoskeleton and sarcomere structures all potentially contribute to *in vivo* functional stiffness. Little is known about how stiffness measured at different structural levels may determine function *in vivo* and how stiffness elements are quantitatively changed in pathophysiology – in particular in acquired disease conditions. Thus, the goal of this investigation was to experimentally characterize *in vitro* stiffness properties (stress/strain) of cardiomyocytes derived from hearts of animals exhibiting diastolic dysfunction *in vivo* in the setting of dietary induction of cardiometabolic disease.

## METHODS & MATERIALS

### Ethical approval

The authors understand the ethical principles under which *The Journal of Physiology* operates, and the work presented in this study complies with the animal ethics checklist as outlined in Instructions to Authors. All experiments conform to the principles and regulations as described by Grundy (2015). Animals (male C57Bl/6J mice) were purchased from the Animal Resource Centre (Perth, WA Australia). All animal experiments were approved by the University of Melbourne (School of Biomedical Sciences) Animal Ethics Committee (Project Ethics Approval 10366) and complied with the guidelines and regulations of the Code of Practice for the Care and Use of Animals for Scientific Purposes (National Health and Medical Research Council Australia). Animals were provided with free access to food and water, were group housed in a temperature-controlled environment 21-23°C with 12 hour light/dark cycles. Animals were euthanised under anaesthesia (single dose of sodium pentobarbital 70 mg/kg i.p) on cardiac excision.

### Animal model of cardiometabolic disease

As a pre-clinical cardiometabolic disease model, the high fat/sugar diet (HFSD) fed rodent recapitulates a common and growing human disease phenotype (obesity, insulin resistance, glucose intolerance) and has been established as a diastolic dysfunction model (Heather *et al*., 2022). Cardiometabolic disease was induced in male C57Bl/6J mice by a high fat/sugar dietary intervention commencing at 8-9 weeks of age. Animals were randomized to experimental treatment groups and were group housed (> 4 animals per cage) in a temperature-controlled environment 21-23°C with 12 hour light/dark cycles. After 1 week transitional feeding period (alternating feed periods), animals were fed a HFSD (43% kcal from fat, 200 g/kg sucrose, SF04-001, Specialty Feeds) or control reference diet (CTRL: 16% kcal from fat, 100 g/kg sucrose, AIN93G, Specialty Feeds) for at least 25 weeks.

### Glucose tolerance testing

At 23-25 weeks of feeding, glucose tolerance testing was performed in mice following 6 hours fasting as previously described (Chandramouli *et al*., 2018). Baseline blood glucose levels were measured using an Accu-Chek glucometer with a blood sample obtained from a needle prick to the tail vein. Glucose (1.5 g/kg body weight) was injected i.p. and blood glucose was measured at 5, 15, 30, 60, 90 and 120 minutes after the glucose injection.

### Transthoracic echocardiography

At 25-26 weeks of feeding, cardiac structure, systolic and diastolic function were evaluated by transthoracic 2-dimensional B- and M-mode and blood flow and tissue doppler echocardiography (GE Vivid 9; 15mHz i13L linear array transducer) as described previously (Chandramouli *et al*., 2018; Curl *et al*., 2018). Briefly, mice were lightly anesthetised with Isoflurane (1.5%; Baxter healthcare, Australia). Parasternal short axis view was used to measure morphological and systolic parameters (interventricular septum IVS, left ventricular posterior wall LVPW, left ventricular internal dimension LVID, fractional shortening FS, ejection fraction EF, heart rate HR). Pulse wave Doppler and tissue Doppler scans were acquired from the apical 4 chamber view to assess LV diastolic function parameters: velocity of early (E) and late (A) mitral inflow and early (e’) and late (a’) diastolic velocity of mitral annulus. From these compound indices of diastolic function E/e’, E/A and e’/a’ were calculated. In addition, the slope of the E wave was used to calculate mitral valve deceleration time. Three consecutive cardiac cycles were sampled for each measurement.

### Cardiomyocyte preparation, structural and functional assessment

A series of protocols were implemented to phenotype cardiomyocyte and sarcomere morphology, and assess functional status in different contraction configurations and challenge settings.

#### Cardiomyocyte isolation, morphology & microfluorimetry

Prior to cardiomyocyte isolation animals were aneasthetised (sodium pentobarbital 70 mg/kg i.p) and hearts excised. HFSD and CTRL left ventricular cardiomyocytes were enzymatically isolated using methods previously described (Howlett *et al*., 2006; Mellor *et al*., 2011) at 30-32 weeks feeding. Briefly, the heart was excised, cannulated via the aorta, and perfused on a Langendorff apparatus with an oxygenated Ca^2+^ free buffer at 37°C for 10 minutes. The buffer was composed of (in mM): 130 NaCl, 5 KCl, 0.33 NaH_2_PO_4_; 1 MgCl_2_, 25 HEPES; 20 Glucose; 3 Na Pyruvate; 1 Na Lactate. The Ca^2+^ free buffer was then supplemented with 0.66mg/mL collagenase (Worthington; Type II), 0.033 mg/mL trypsin and 0.05 mM CaCl_2_ and perfusion continued for a further 10 minutes. The heart was removed from the Langendorff apparatus, and the left ventricle was segmented into small pieces in a high potassium solution composed of (in mM): 30 KCl, 90 KOH, 30 KH_2_PO_4_, 3 MgSO_4_, 50 glutamic acid, 20 taurine, 0.5 EGTA, 10 glucose, and 10 HEPES. Single left ventricular cardiomyocytes were isolated via gentle trituration. Viable cardiomyocytes were resuspended in HEPES-Krebs buffer (in mM); 146.2 NaCl, 4.69 KCl, 0.35 NaH_2_PO_4_H_2_O, 1.05 MgSO_4_7H_2_O, 10 HEPES, 11 glucose containing 1.8 mM Ca^2+^ and were stored on a rocking platform at room temperature. From each heart cell dimensions were collected from at least 50 rod shaped cardiomyocytes using an inverted light microscope (Nikon) and calibrated eye piece as previously described (Curl *et al*., 2018). Cardiomyocyte volume was calculated by inserting experimentally determined myocyte length and width values into the formula: volume = length x width x depth x multiplier 0.54 (elliptical area of the rectangle defined by width & length, (Satoh *et al*., 1996)). Cardiomyocyte width: depth ratio was assumed to be 1.44 (Natali *et al*., 2001). Cardiomyocyte cross-sectional area (CSA) could be calculated from measured width scaled for 2-dimensional geometry (CSA = W [W/1.44] x 0.54 µm^2^). An aliquot of isolated cardiomyocyte suspension was pipetted into a cell chamber mounted on the stage of an inverted fluorescence microscope (Motic AE31) and superfused with HEPES-Krebs solution with 2 mM Ca^2+^, 37°C at a flow rate of 2 mL/min. Cardiomyocyte sarcomere length and Ca^2+^ transients were simultaneously recorded during isotonic contractions. Myocyte force development was evaluated in the same cells with transducer attachment, using four protocols described below.

#### Characterizing cardiomyocyte diastolic function: non-loaded shortening measurements (isotonic), loaded force measurements (auxotonic), stretch responses and pacing challenge

Cardiomyocyte performance was evaluated in several paced conditions: designated ‘non-loaded’ (isotonic), ‘loaded’ (auxotonic after-loaded), and ‘stretched’(auxotonic pre-loaded). The non-loaded state refers to the absence of externally applied load (noting that sarcomeric and cytoskeletal structures do constitute an underlying internal load).

Firstly, cardiomyocyte non-loaded (isotonic) sarcomere contractions (IonOptix SarcLen, Fast Fourier Transform, FFT) and Ca^2+^ transients (Fura2-AM, 5 μM microfluorimetry F340:380 nm ratio) were simultaneously measured at 2 Hz, using a microscope diaphragm to define a rectangular intra-myocyte region of interest as depicted. During contraction cycles, cardiomyocyte sarcomere length was monitored within an intracellular region of interest defined by an adjustable diaphragm mounted in the microscope optical tube (**Fig 4A**).

Secondly, to obtain loaded (auxotonic) contraction data, each cardiomyocyte was attached to a pair of glass fibres coated with a bio-adhesive (MyoTak, IonOptix, USA). Attachment sites were positioned to be outside the sarcomere tracking region, closer to myocyte longitudinal ends. One glass fibre was connected to a force transducer (fibre stiffness 20 N/m, IonOptix, USA), and the other to a piezo-motor (Nano-drive, Mad City Labs Inc. WI, USA) to adjust cardiomyocyte length (**Fig. 4A**). With successful attachments, cardiomyocytes were lifted from the coverslip using motorised micromanipulators (MC1000E, Siskiyou Corporation, USA) attached to both the force transducer and the piezo-motor. Baseline loaded contractions at 2 Hz were recorded. Sarcomere length changes were measured using FFT as described above. In addition, with glass fibre attachment, tracking of mean sarcomere shortening within a myocyte segment delineated by the glass fibre edge positions could be performed in parallel (**Fig4B-C**).

Thirdly, force was recorded during a standardized stretch protocol of loaded cardiomyocytes to measure the effect of increasing diastolic cardiomyocyte/sarcomere basal length during pacing. A stress-length relation was constructed for each myocyte (where stress = force/cross-sectional area). Prior to the commencement of the myocyte stretch measurements a protocol to calibrate the stretch step was implemented. Four consecutive 0.5 V input signals were delivered to the piezo-motor to determine a calibration constant to define the percentage cell stretch for a given voltage (**Fig. 4B**). For each myocyte a *Calibration Constant* was calculated as: *myocyte segment length change per piezo-motor mV / sarcomere count in segment* [where *myocyte sarcomere count = myocyte segment length (µm) / average diastolic sarcomere length (µm)*]. To track the glass fibre position a real-time contrast detection algorithm (IonOptix - EdgeLength) was focused on the internal edge of each of the glass fibres (**Fig. 4**) (Peyronnet *et al*., 2017). This enabled reproducible and comparable stretch steps to be applied to different cardiomyocyte preparations (**Fig. 4B-C**). Prior to the ‘stress-length’ protocol, a cell-specific calibration constant was applied to yield a standardised stretch for each cardiomyocyte evaluated. The stress-length protocol involved progressive stretch steps totaling an approximate 30% cell stretch (**Fig. 4B-C**).

Lastly, the pacing was increased to 4 Hz, a rate adaptation period allowed (2 min) and the stress-length protocol was repeated. Sarcomere length, force development and intracellular Ca^2+^ transients were simultaneously measured. An average of at least 10 twitches or transients were used to calculate sarcomere, Ca^2+^ and force parameters at a given stretch length. In summary, the following parameters were measured: diastolic stress (mN/mm^2^), diastolic stress-length relation [(mN/mm^2^) / (%^-^ length change)], diastolic stress-sarcomere length relations ((mN/mm^2^) / µm), systolic : diastolic stress-sarcomere relation (Frank Starling Gain Index (Bollensdorff *et al*., 2011)), systolic : diastolic stress-length relation, systolic stress amplitude (mN/mm^2^), sarcomere shortening (% diastolic SL), maximum sarcomere shortening and lengthening velocity (µm/s). In addition, Ca^2+^ parameters were measured: diastolic Ca^2+^, systolic Ca^2+^ and time constant of decay of the Ca^2+^ transient (tau).

### Statistical analyses

Data are presented as mean ± standard deviation and statistical analysis was performed using GraphPad Prism V9.5.1. Cardiomyocyte stress-length and stress-SL relations were modelled with simple linear regression. For comparison between two groups an unpaired student t-test was used. For assessment between two groups at multiple points a one-way ANOVA with Bonferroni multiple comparisons post-hoc test was used. Two-way ANOVA with Šídák’s multiple comparisons post-hoc test was used for the evaluation of data groups with two independent variables. Pearson Correlation analyses were used to assess statistical significance of fitted linear regression lines. A p-value of <0.05 was considered statistically significant. Analyses were performed using a blinded protocol.

## RESULTS

### Systemic indicators of cardiometabolic disease and *in vivo* diastolic function in HFSD mice

High fat/sugar diet-fed mice exhibited obesity and glucose intolerance. Relative to CTRL, the body weight of HFSD mice was significantly elevated within 5 weeks of dietary intervention increasing over the study duration by 50% relative to CTRL (**Fig. 1A**, CTRL: 28.1±2.0 vs. HFSD: 44.3±1.9 g). After 23-24 weeks diet treatment, glucose tolerance testing of a subset of the HFSD cohort was performed. Glucose intolerance in response to intraperitoneal glucose administration was evidenced by increased area under the curve, consistent with literature reporting insulin resistance in mice fed a similar diet (**Fig. 1B**).

**Fig. 1.**
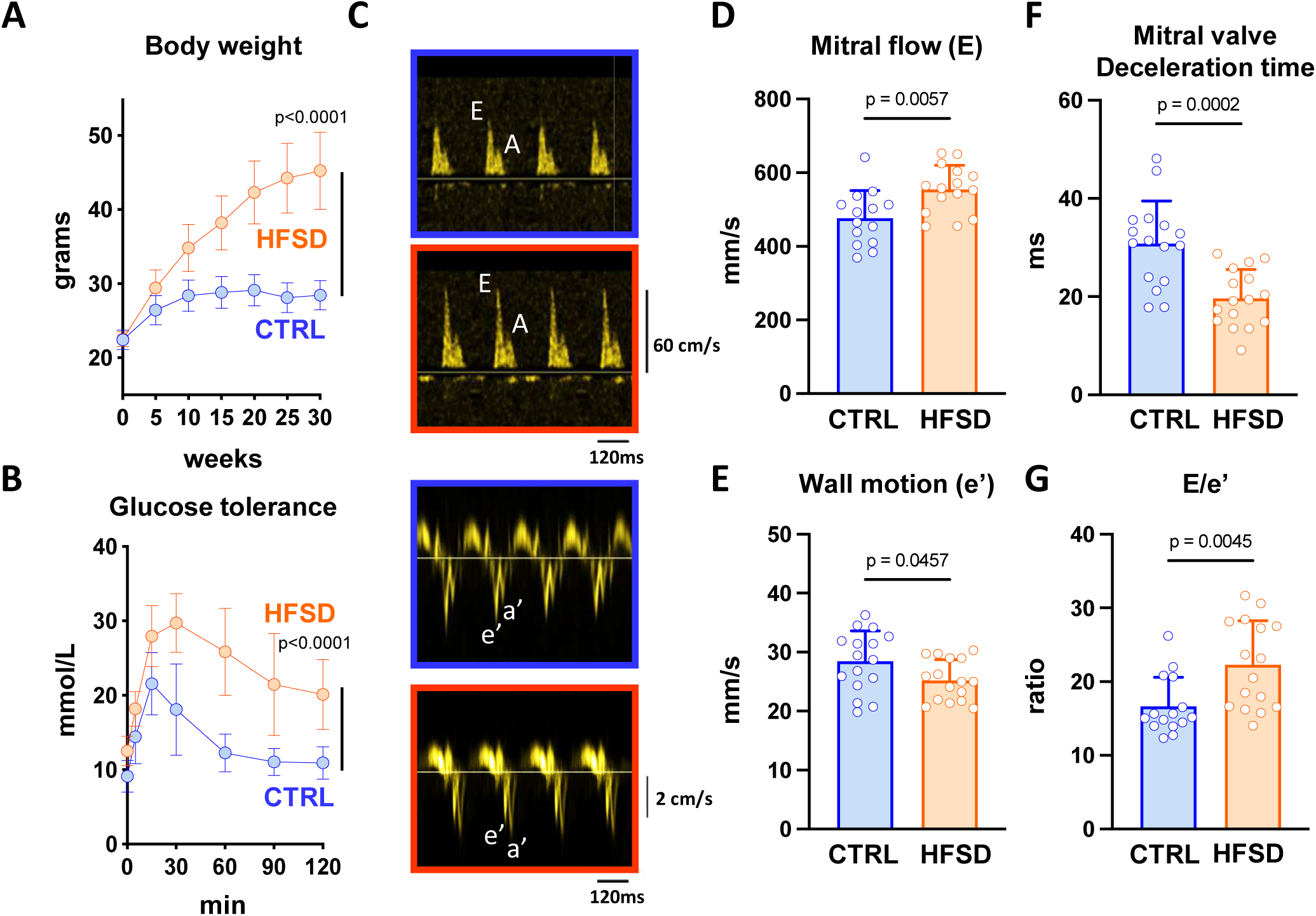
Systemic indicators of cardiometabolic disease and *in vivo* diastolic function in high-fat/sugar diet (HFSD) vs control (CTRL) mice. **A.** Body weight elevation at 5 week intervals up to 30 weeks of dietary treatment (last timepoint full cohort intact. N = 16 – 17 mice per group). **B.** Glucose tolerance testing at 23-24 weeks post dietary intervention; N = 6 mice per group). **C.** Echocardiography exemplar blood flow and tissue doppler images from which diastolic indices were derived for CTRL and HFSD mice. Panels with orange outline: HFSD mitral blood flow & annular wall motion. Panels with blue outline: CTRL mitral blood flow & annular wall motion. (N = 14-16 mice per group) **D.** Early diastolic mitral annular blood flow (E) was increased in HFSD. **E.** Early diastolic mitral annular wall velocity (e’) was decreased in HFSD. **F.** Deceleration time of early diastolic mitral annular blood flow was decreased in HFSD. **G.** Ratio of early diastolic blood flow to wall velocity (E/e’) was increased in HFSD. Time-course data analysed using two-way repeated measures ANOVA with Sidak post-hoc test with p-value for ‘time x diet’ effect presented. Comparisons between two treatment groups performed using student’s t-test. Data presented as mean ± standard deviation.

After confirming these key cardiometabolic disease characteristics in the HFSD mice, echocardiographic functional evaluation was undertaken. Focusing on diastolic function evaluation (**Fig. 1C-G**), there was higher early mitral inflow (**Fig. 1D, E** wave, CTRL: 472±74, HFSD: 569±65 mm/s) and reduced velocity of wall motion (**Fig. 1E**, e’ wave, CTRL: 28.5±5.1, HFSD: 25.2±3.5 mm/s) in HFSD myocardium. Deceleration time of mitral inflow was shortened (**Fig. 1F**, CTRL: 30.8±8.7, HFSD: 19.7±5.9 ms) and E/e’ ratio increased (**Fig. 1G**, CTRL: 16.7±3.9, HFSD: 22.3±6.0), all observations consistent with reduced tissue compliance and diastolic dysfunction. No significant differences in systolic performance parameters were observed (**Table. 1**), confirming the primary *in vivo* phenotype of diastolic dysfunction. The next step was to evaluate the cardiomyocyte involvement in dysfunction.

**Table 1.** Echocardiographic parameters in CTRL and HFSD mice. *LVIDd* left ventricular internal diameter at end diastole, *IVSd* interventricular septum diameter at end diastole, *LVPWd* left ventricular posterior wall thickness at end diastole, *RWT* relative wall thickness, *E wave* mitral valve blood flow velocity during the early ventricular filling phase, *A wave* mitral valve blood flow velocity during atrial contraction, *e’ wave* mitral annulus tissue motion velocity during the early ventricular filling phase, *a’ wave* mitral annulus tissue motion velocity during atrial contraction, *MV* mitral valve. Comparisons between two treatment groups performed using Students’ unpaired t-test (N = 9-15 mice per group). Data are presented as mean ± standard deviation.

| **Table 1. Echocardiographic parameters for CTRL and HFSD mice** |  |  |  |
| --- | --- | --- | --- |
|  | **CTRL** | **HFSD** | **P-value** |
| Heart rate (bpm) | 515±74 | 551±29 | 0.174 |
| LV mass (mg) | 125±27 | 131±21 | 0.314 |
| LVIDd (mm) | 3.60±0.17 | 4.16±0.47 | 0.004 |
| IVSd (mm) | 0.84±0.06 | 0.80±0.10 | 0.349 |
| LVPWd (mm) | 0.94±0.03 | 0.94±0.10 | 0.830 |
| RWT (ratio) | 0.49±0.06 | 0.43±0.09 | 0.079 |
| Ejection Fraction (%) | 67.8±3.7 | 64.9±3.2 | 0.093 |
| Fractional Shortening (%) | 31.8±3.6 | 30.6±2.1 | 0.400 |
| E wave (mm/s) | 472±74 | 569±65 | 0.003 |
| A wave (mm/s) | 348±99 | 400±116 | 0.190 |
| e' wave (mm/s) | 28.5±5.1 | 25.2±3.5 | 0.046 |
| a' wave (mm/s) | 23.5±7.8 | 23.5±6.4 | 0.995 |
| E/A | 1.51±0.41 | 1.47±0.37 | 0.095 |
| e'/a' | 1.33±0.44 | 1.12±0.39 | 0.171 |
| E/e' | 16.7±3.9 | 22.3±6.0 | 0.005 |
| MV Deceleration Time (ms) | 30.8±8.7 | 19.7±5.9 | <0.001 |

### Non-loaded sarcomere and Ca^2+^ dynamics in CTRL and HFSD cardiomyocytes

Single isolated cardiomyocytes were prepared from hearts of a subset of mice which underwent echocardiographic analyses. For intact, non-loaded cardiomyocytes sarcomere shortening and lengthening cycles were similar for CTRL and HFSD groups (**Fig. 2A**). Diastolic sarcomere length (SL) (**Fig. 2B**, CTRL: 1.71±0.04, HFSD: 1.72±0.04 µm) and extent of sarcomere shortening (**Fig. 2C**, CTRL: 3.79±2.24, HFSD: 3.94±3.15 % diastolic SL) were not significantly different between groups. Examination of sarcomere contraction kinetics (**Fig. 2D** exemplar trace) showed that the maximum rate of lengthening (**Fig. 2E**, CTRL: 1.08±0.85, HFSD: 1.29±1.07 μm/s) and shortening (**Fig. 2F**, CTRL: −2.32±1.44, HFSD: −2.57±2.01 μm/s) did not significantly differ between CTRL and HFSD groups. Collectively these findings indicate that in the non-loaded setting no differences in CTRL and HFSD cardiomyocyte systolic and diastolic non-loaded performance were detected, a finding consistent with other studies investigating non-loaded cardiomyocyte performance in diet-induced cardiometabolic disease including our previous studies in fructose-fed mice (Mellor *et al*., 2012; Llano-Diez *et al*., 2016; West *et al*., 2019; Wang *et al*., 2020).

**Fig. 2.**
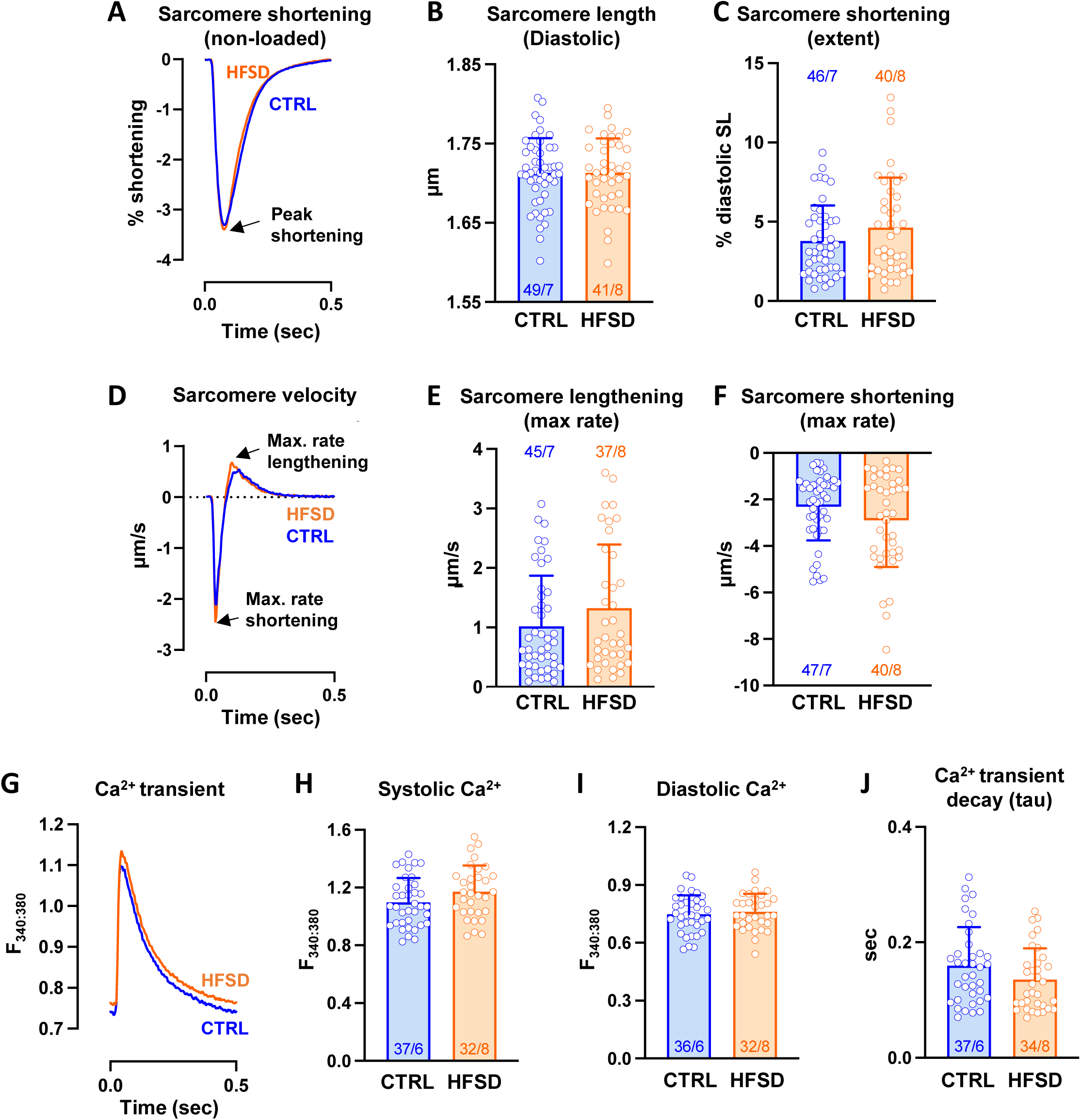
Non-loaded sarcomere and Ca^2+^ dynamics in CTRL and HFSD cardiomyocytes. **A.** Mean sarcomere contraction cycles presented as a percentage of diastolic sarcomere length for CTRL and HFSD cardiomyocytes **B-C.** Mean diastolic sarcomere length and extent of sarcomere shortening for CTRL and HFSD cardiomyocytes. **D.** Mean sarcomere shortening and lengthening rate records (time derivative of sarcomere contraction-relaxation cycle) for CTRL and HFSD cardiomyocytes. The nadir and zenith represent maximal shortening and lengthening rates respectively. **E-F.** Maximal rate of lengthening and shortening rates for CTRL and HFSD cardiomyocytes. **G.** Mean Ca**^2+^** transient records for CTRL and HFSD cardiomyocytes. **H-J.** Mean Ca**^2+^** levels (systolic and diastolic) and time constant of transient decay (*tau*) in HFSD and CTRL cardiomyocytes. Mean data computed from average of 20 cycles per cardiomyocyte to derive group mean (N = 6-8 mice per group, n = 32-49 cells per group). Comparisons between two treatment groups performed using student’s t-test (all p>0.05, ns). Data presented as mean ± standard deviation.

Next, Ca^2+^ parameters were compared with cardiomyocyte sarcomere lengthening performance during relaxation and diastole (**Fig. 2G**, exemplar traces). No significant difference was evident between dietary treatment groups in the systolic Ca^2+^ (**Fig. 2H**, CTRL: 1.098±0.169, HFSD: 1.171±0.182 F_340:380_), diastolic Ca^2+^ (**Fig. 2I**, CTRL: 0.749±0.097, HFSD: 0.761±0.094 F_340:380_) or time constant (tau) of Ca^2+^ transient decay (**Fig.2J**, CTRL: 0.160±0.067, HFSD: 0.137±0.054 sec). Thus, the significant diastolic dysfunction measured *in vivo* for the HFSD group was not observed in a non-loaded *in vitro* setting. To reconcile the impaired echocardiographic heart performance with absence of dysfunction in non-loaded contracting single cardiomyocytes, the next strategy was to assess single cardiomyocyte response to loading. A common feature of cardiometabolic disease is elevation in after-load – i.e. elevated systemic resistance potentially impeding myocardial pump function. To simulate this *in vitro*, mechanically loaded cardiomyocyte performance was examined.

### Loaded cardiomyocyte sarcomere shortening and lengthening

Immediately after non-loaded cardiomyocyte recordings, glass fibres were attached to cardiomyocyte longitudinally separated positions to permit auxotonic, loaded contraction measurements. For cells successfully attached, the mean difference in sarcomere length measured pre- and post-conversion to loaded mode was <0.02% diastolic length, indicating non-loaded and loaded sarcomere initial lengths were well matched. For attached cardiomyocytes, a comparison of sarcomere shortening behavior in the loaded state could be made (using FFT) to directly match with data obtained non-loaded state.

In the first instance, data from all myocytes were pooled to investigate the general effects of load regardless of treatment group. With loading, for all myocytes (both treatment groups), extent of sarcomere shortening was reduced and shortening and lengthening kinetics slowed when compared with non-loaded performance (**Fig. 3A**). Mean data showed that in response to loading, sarcomere shortening was reduced by ∼34% (**Fig. 3B**, non-loaded: 0.075±0.042, loaded: 0.050±0.031 µm). Maximum rate of shortening was decreased by 58% (non-loaded: −2.85±1.40, loaded: −1.20±0.89 µm/s; not shown), and maximum rate of lengthening by 66% (**Fig. 3C**, non-loaded: 1.46±1.03, loaded: 0.52±0.50 µm/s). The magnitude of the change in extent of shortening and maximum rate of lengthening with loading were positively correlated (**Fig. 3D**) suggesting the possibility of common underlying mechanisms.

**Fig. 3.**
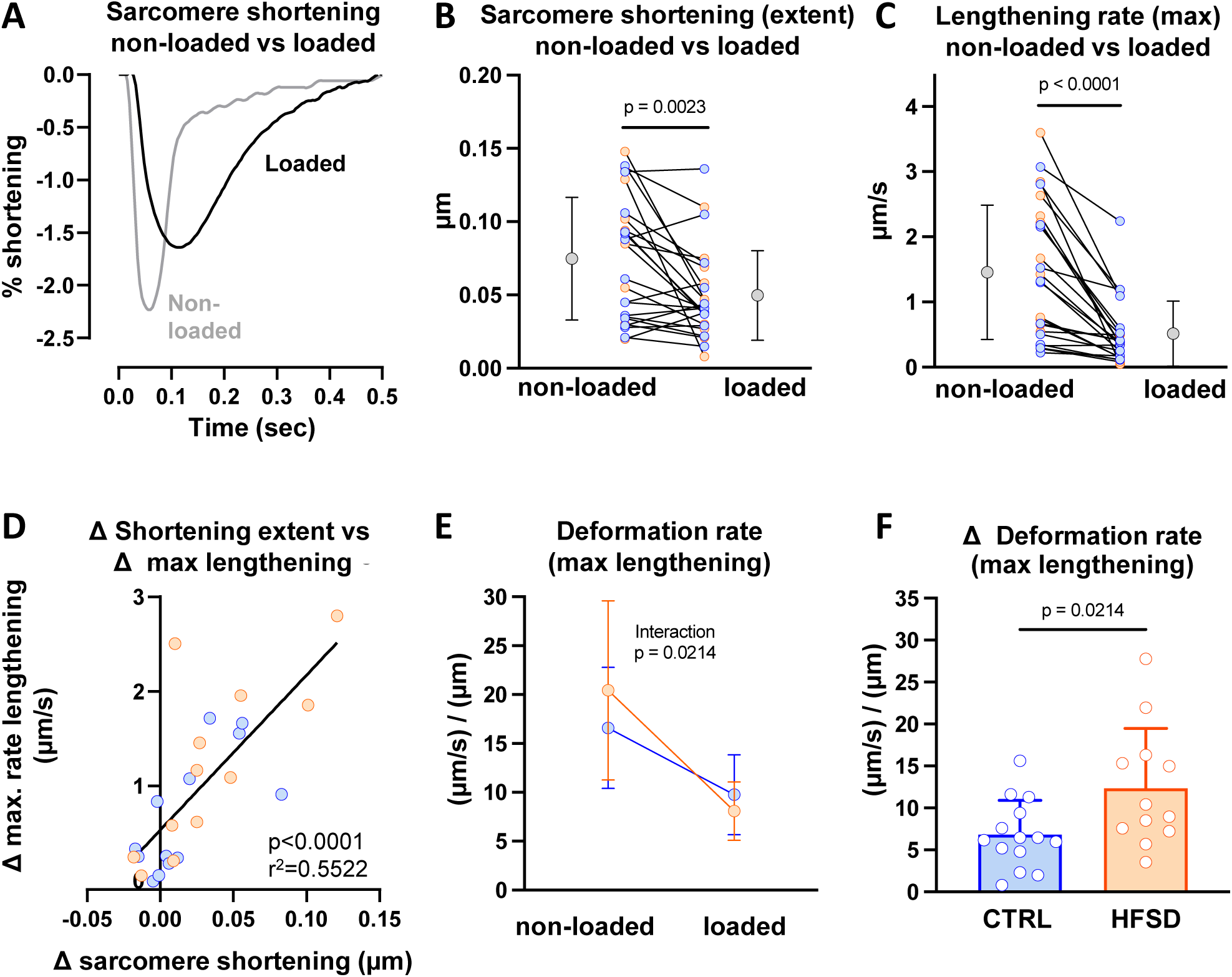
Sarcomere shortening and lengthening in non-loaded and loaded pacing states. **A.** Exemplar non-loaded and loaded sarcomere contraction cycles for the same cardiomyocyte (CTRL). **B-C.** Extent of sarcomere shortening and maximum rate of lengthening, measured for each cardiomyocyte in non-loaded and loaded state for CTRL and HFSD groups combined (N = 9 mice, n = 25 cells). Mean data for CTRL and HFSD are presented as grey points to the left (non-loaded) and right (loaded) of individual cardiomyocyte data. **D.** Correlation of loading associated sarcomere shortening change and maximum rate of lengthening change for CTRL and HFSD combined. **E.** Mean loading effect on maximum deformation rate during lengthening ((µm/s) / µm; maximum rate of lengthening (µm/s) normalized by shortening extent (µm)) for CTRL vs HFSD. (CTRL: N = 5 mice, n = 14 cells; HFSD: N = 4 mice, n = 12 cells). Significant ANOVA interaction effect (diet treatment x load state). **F.** Mean decrement in deformation rate during lengthening ((µm/s) / µm). Comparison of cardiomyocyte responses in different loading conditions analysed by paired t-test. Comparison of cardiomyocytes from different treatment groups analysed by unpaired t-test. A 2-way ANOVA with repeated measures used for analysis of two independent variables. Data fitted with linear regression model and calculation of Pearson’s correlation coefficient. Data presented as mean ± standard deviation.

To assess cardiomyocyte rate kinetics in non-loaded and loaded cardiomyocytes independent of shortening extent, the maximum rates of lengthening and shortening were normalised by shortening amplitude to derive ‘deformation rate’ - i.e. rate of length change, relative to extent of shortening ((µm x sec^-1^)/µm = 1/sec). The effect of loading on deformation rate was accentuated in HFSD group (significant ANOVA diet x load interaction effect, **Fig. 3E**). A selective HFSD effect was revealed in the differential between matched ‘loaded’ and ‘non-loaded’ lengthening cardiomyocytes. The decrement in deformation rate (lengthening) for CTRL and HFSD is shown in **Fig. 3F**, and is accentuated in HFSD in response to loading (CTRL: 6.84±4.08, HFSD: 12.35±7.15 1/s). Treatment differences in deformation rate were not evident during the shortening phase (data not shown). These findings indicated an accentuated influence of load, specific to the lengthening (i.e. diastolic) phase in HFSD cardiomyocytes and could reflect impaired subcellular relaxation processes specific to the HFSD myocardium. Based on these data identifying different CTRL and HFSD origin cardiomyocyte responses in loaded state, we next investigated the effect of stretch applied to cardiomyocytes to simulate *in vivo* ‘pre-load’ conditions.

### Cardiomyocyte force development: measuring strain & stress during progressive cardiomyocyte stretch

A hallmark of diastolic dysfunction is an elevated diastolic pressure-volume relationship which describes the increased resistance of the myocardial wall to chamber filling (Aurigemma *et al*., 2006). We sought to design a single cardiomyocyte version of this relationship (Fowler *et al*., 2015), by applying standardized stretch steps to cardiomyocytes, defining the steps as a % of non-stretched state (i.e. ‘strain’). Diastolic force was measured simultaneously during length stretch steps. The force measured at every strain increment was normalized by cross sectional area (CSA) to produce a measure of ‘stress’ (i.e. force/CSA, mN/mm^2^). Thus a sarcomere ‘stress/strain’ relation (i.e. force-length) was constructed for each myocyte. The protocols employed to generate stress-strain are depicted in **Fig. 4** and described in detail in the Methods. **Fig. 4** shows the stretch standardization approach used to enable reproducible application of strain (converting voltage signals to linear displacement) as a % length change (via individual myocyte calibration constant calculation) to allow comparisons between different cardiomyocytes. The size of each step change in length on sarcomere length was measured (FFT) and confirmed by tracking the intra-cardiomyocyte segment between the glass fibres (**Fig. 4**). In some cases a direct sarcomere FFT measure was not microscopically resolvable due to myocyte contraction responses which partially positioned some area of the myocyte outside the optimal focal plane. In these situations it was always possible to calculate segment length change. For the example shown, stretch steps were determined to be 2.75µm, and converted to % sarcomere length (i.e. strain) using sarcomere count within intracellular region of interest area (**Fig. 4C**). A comparison of the variability of cell stretches performed using a uniform (pre-set) stretch constant and the individually derived stretch calibration constant demonstrates that reproducibility is optimized with usage of the individual stretch calibration constant data (**Fig. 4D**). Force, sarcomere length and % stretch were measured simultaneously (**Fig. 4**).

**Fig. 4.**
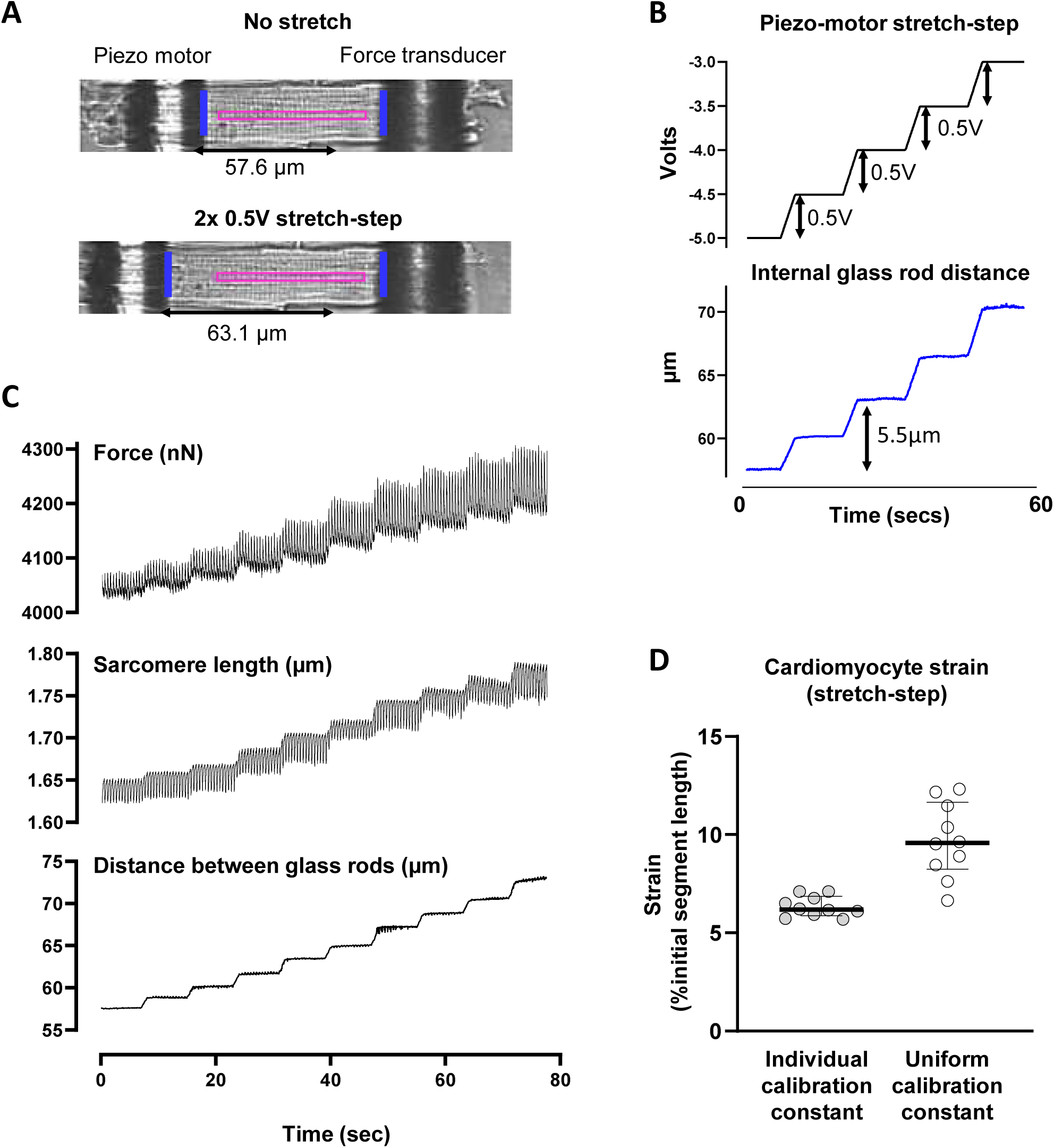
Cardiomyocyte standardized stretch protocol. **A.** Image of a left ventricular cardiomyocyte longitudinally attached to glass rods. Edge-tracking used to measure inter-rod distance (marked as blue bars) for cardiomyocyte segment analyses. The delineated region of interest (marked pink) represents the area from which Fast Fourier Transform analysis of sarcomere striations was performed to derive sarcomere-specific data. **B.** Exemplar traces depict the change in glass rod internal distance in response to successive 0.5V piezo motor inputs used for calibration of stretch protocol for each individual cardiomyocyte to allow normalized comparisons. **C.** Measuring stress and strain during progressive cardiomyocyte stretch. Exemplar force, sarcomere length and rod edge-tracker traces during the first stretch of a stress-length protocol. **D.** Comparison of variability of cell stretches performed using a uniform (preset) stretch constant and the individually derived stretch calibration constant. Strain was measured as the %change in segment length relative to the initial segment length. Reproducibility is optimized using the individual stretch calibration constant. Data shown as median ± interquartile range.

### Quantifying diastolic stiffness in CTRL vs HFSD cardiomyocytes using strain normalization methods

Superimposed exemplar stress recordings are shown in **Fig. 5A** for CTRL and HFSD cardiomyocytes, captured during a series of progressive stretches during a 100 sec period. The dashed (broken) lines are aligned with the diastolic stress step increments for each of the 2 myocytes (using the data points collected prior to the transition to a new step) throughout the series of progressive stretches. A robust increase in systolic force with stretch is also evident, indicative of length-dependent activation, the cardiomyocyte basis for the Frank-Starling mechanism.

**Fig. 5.**
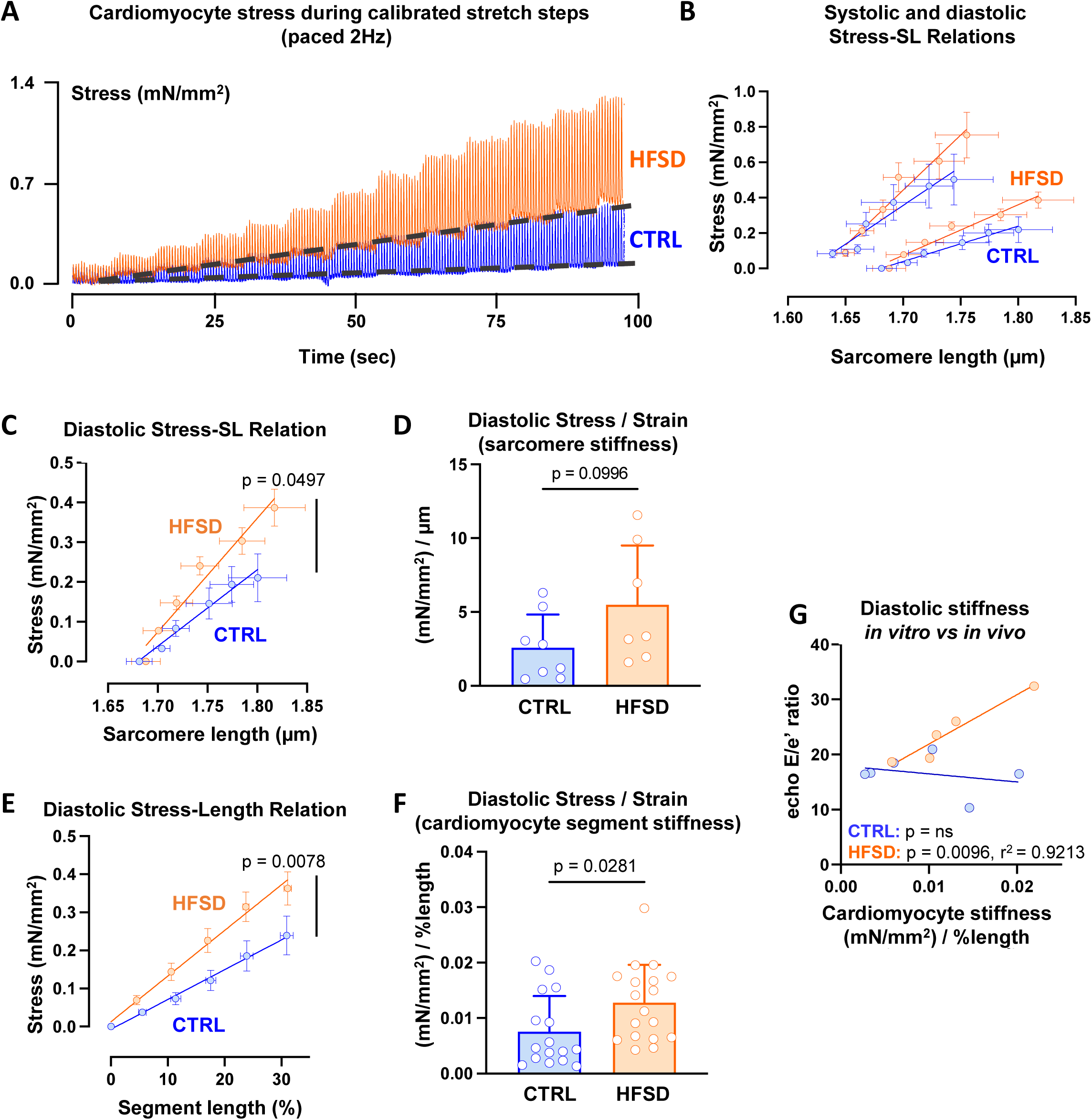
Quantifying diastolic stiffness in CTRL vs HFSD cardiomyocytes using strain normalization methods. **A.** Exemplar stress recordings of CTRL and HFSD cardiomyocytes undergoing ‘stress-length’ protocol (2Hz paced). Diastolic baseline stress shifts indicated by broken line. **B.** Diastolic (pair of lower lines) and systolic (pair of upper lines) sarcomere mean stress-sarcomere length (SL) relations in CTRL cardiomyocytes (N = 5 mice, n = 9 cells). **C.** Diastolic mean stress-sarcomere length relation (linear regression fit) for HFSD and CTRL. **D.** Diastolic stress (mN/mm^2^) normalized relative to sarcomere stretch (µm) yields sarcomere stress/strain = stiffness [(mN/mm^2^) /strain (µm)] for HFSD and CTRL. **E.** Diastolic mean cardiomyocyte segment stress-length relation (linear regression fit) for HFSD and CTRL.. **F.** Diastolic stress (mN/mm^2^) normalized relative to segment stretch (% length) yields cardiomyocyte segment stress/strain = stiffness [(mN/mm^2^) /strain (% length)] for HFSD & CTRL. **G.** Correlation (linear regression fit) of *in vitro* cardiomyocyte segment diastolic stiffness (stress/strain) with *in vivo* diastolic dysfunction E/e’ (blood flow/wall motion) in CTRL and HFSD. Cardiomyocyte data averaged per heart. Comparisons between two groups performed using student’s t-test. Data fitted with linear regression model and calculation of Pearson’s correlation coefficient, presented as mean ± standard deviation (D, F) and mean ± standard error of the mean (B, C, E). For **C. & D.** (CTRL: N=4 mice, n=8 cells; HFSD: N=2 mice, n=7 cells). For **E.&F.** (CTRL: N=6 mice, n = 16 cells; HFSD: N=6 mice, n=18 cells).

In this experimental setting, stiffness could be visualized as the gradient of the linear plot derived from the cardiomyocyte diastolic stress-length relation. Our stretch protocol methodology provided two approaches for stiffness calculation (as per Methods). A first approach used FFT measurement of sarcomere length to track cardiomyocyte stretch and was plotted according to the standardized stretch steps (in µm). An alternative approach used percent length change calculated as the length of the cardiomyocyte segment between the glass attachment fibres divided by the initial distance between the fibres prior to stretch (in % initial length).

Applying the first approach, a linear regression was performed to fit the data from which a slope value could be derived. **Fig. 5B** depicts the average diastolic and systolic stress-sarcomere length relations for CTRL cardiomyocytes only. To determine cardiomyocyte diastolic stiffness using FFT-computed sarcomere length values, the gradient of the mean slope of the diastolic stress sarcomere-length relation for each treatment group was calculated, using mean sarcomere length change (μm) as the denominator. This gradient was significantly higher in HFSD myocytes compared to CTRL (**Fig. 5C**). The individual slope gradients were used to calculate the mean diastolic stress / strain gradient (**Fig. 5D**, CTRL: 2.58±2.25, HFSD: 5.50±4.01 (mN/mm^2^) / µm, p = 0.0996) where there was a trend for a 2.3 fold increase in stiffness of HFSD cardiomyocytes. The statistical power of this analysis was limited due to the difficulty in achieving resolvable diastolic sarcomere lengths by FFT for all myocytes.

Applying the second approach, it was possible to construct stress-segment length (%) relations for all myocytes from which stress data could be obtained. In **Fig. 5E**, the stress-segment length data for HFSD and CTRL are shown, with a robustly significant difference detected in mean gradient values. In **Fig. 5F** the diastolic stress/strain values calculated for the myocytes from the two treatment groups using this segment approach are directly compared. The HFSD cardiomyocyte stiffness index derived using this approach was increased by 70% compared with CTRL (**Fig. 5F**, CTRL: 0.0075±0.0064, HFSD: 0.0128±0.0068 (mN/mm^2^) / % length). There was no significant difference in ‘Frank Starling Gain index’ (**Table. 2,** CTRL: 2.40±1.06, HFSD: 2.77±1.45, (mN/mm^2^) / µm) or the ratio of systolic to diastolic stress segment length relations (**Table. 2,** CTRL: 1.62±0.25, HFSD: 1.73±0.50, (mN/mm^2^) / % length) indicating that cardiomyocyte length-dependent activation is preserved (Bollensdorff *et al*., 2011; Najafi *et al*., 2016).

**Table 2.**
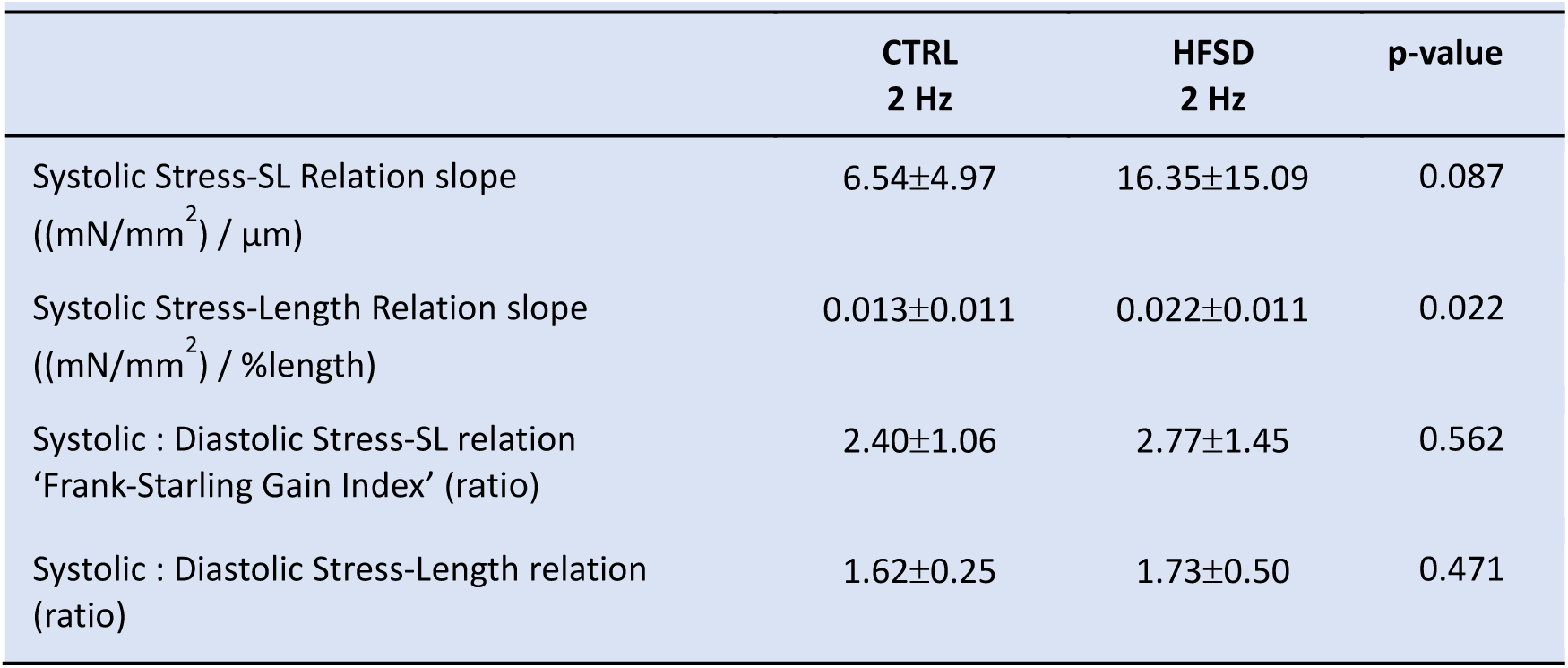
Stress-length relation mean values for CTRL and HFSD mouse cardiomyocytes. Comparisons between two treatment groups performed using Students’ unpaired t-test (CTRL: N = 4-6 mice, n = 9-17 cells cells; HFSD: N = 2-6 mice, n = 7-19 cells). Data are presented as mean ± standard deviation.

To examine the relationship between intrinsic *in vitro* cardiomyocyte stiffness and *in vivo* diastolic dysfunction, diastolic stiffness (slope of the diastolic stress segment-length relation) averaged per animal was plotted against the E/e’ index of diastolic dysfunction. There was a significant correlation between cardiomyocyte stiffness and E/e’ in HFSD and no evidence of a relationship for the CTRL group (**Fig. 5G** CTRL: p = ns, HFSD: p = 0.0096, r^2^ = 0.9213). These data support the interpretation that intrinsic cardiomyocyte stiffness represents a definable and inherent component of diastolic dysfunction occurring *in vivo* in cardiometabolic disease.

### Load dependence of Ca^2+^ transient decay in CTRL and HFSD cardiomyocytes

To further understand the cardiomyocyte functional elements comprising increased intrinsic stiffness in cardiopathology, the relationship between Ca^2+^ transient features and myofilament deactivation in response to progressive stretch was evaluated. In the intact cardiomyocyte model these electro-mechanical coupling processes are functional for interrogation (in contrast to the disrupted cytosolic environment which characterizes permeabilized myocyte preparations). The Ca^2+^ transient decay features were compared for non-loaded and loaded basal states and also in response to progressive stretch.

In CTRL cardiomyocytes there was no difference in the time constant of Ca^2+^ transient decay (tau, 1/e) between loaded and stretched state (20% increase in length, **Fig. 6A**. loaded 1.18±0.48; stretched 1.12±0.35 fold change vs non-loaded reference 1.00±0.46). In HFSD cardiomyocytes the time constant of Ca^2+^ transient decay was increased in stretched cardiomyocytes relative to non-loaded (**Fig. 6A**. non-loaded, 1.00±0.42; loaded 1.22±0.50; stretched 1.42±0.65 fold change). Loading prolonged time to peak Ca^2+^ transient in both CTRL and HFSD cardiomyocytes (**Fig. 6B**. CTRL: non-loaded 1.00±0.25; loaded 1.49±0.38; stretched 1.46±0.44, HFSD: non-loaded 1.00±0.41; loaded 1.36±0.41; stretched 1.21±0.34 foldchange). The effects can be visualized in **Fig. 6C-D** where the mono-exponential decay of exemplar Ca^2+^ transients is depicted. The shallower decay of the Ca^2+^ transient in the stretched state (**Fig 6D**, HFSD) was apparent whereas the loaded and non-loaded decay traces were virtually overlayed for CTRL cardiomyocytes (**Fig. 6C**). These data suggest that stretch may enhance a mechanism by which the myofilaments bind Ca^2+^ for a prolonged duration during relaxation thus reducing Ca^2+^ availability for extrusion from the cytosol.

**Fig. 6.**
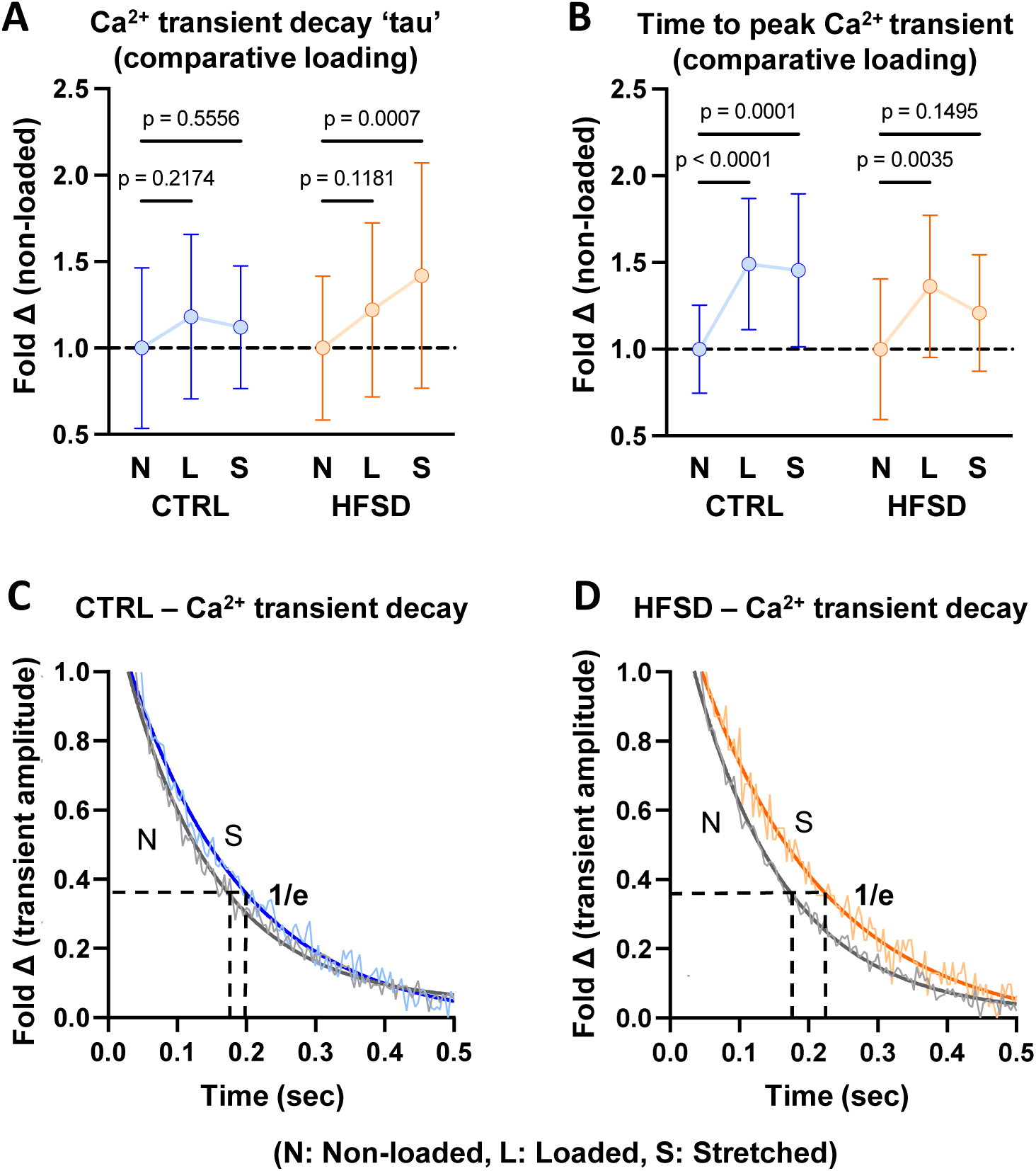
Load dependence of cardiomyocyte Ca^2+^ transient decay in CTRL and HFSD cardiomyocytes. **A-B.** Time constant of Ca^2+^ transient decay (tau) and time to peak Ca^2+^ transient in cardiomyocytes with different load conditions: loaded (L) & stretched (S) shown as fold change relative to non-loaded (N) for CTRL and HFSD. **C-D.** Exemplar Ca^2+^ transient decay records, with a mono-exponential decay function fit, in non-loaded and stretched CTRL and HFSD cardiomyocytes. Horizontal black broken lines intersect with mono-exponential fit at 36.8% (1/e) transient amplitude. Vertical broken lines from intersection points indicate the time constant of Ca^2+^ transient decay (tau). For analysis of two independent variables a 2-way ANOVA with repeated measures used (CTRL: N = 5 mice, n = 13 cells; HFSD: N = 4 mice, n = 12 cells). Data are presented as mean ± standard deviation.

### Pacing-induced [Ca^2+^]_i_ increase to examine potential for a cross-bridge derived stiffness mechanism

A further mechanistic study to probe the potential relationship between Ca^2+^ and stiffness involved perturbation of cytosolic Ca^2+^ cycling time. Elevating stimulation frequency is a recognized physiologic strategy to increase diastolic Ca^2+^ levels and to apply increased cardiomyocyte workload. In response to frequency elevation (2 Hz to 4 Hz) cardiomyocyte diastolic Ca^2+^ levels increased in both CTRL and HFSD (**Fig. 7A & B**) groups consistent with previous studies (Varian & Janssen, 2007; Eisner *et al*., 2020). In the HFSD cardiomyocytes this effect was markedly diminished and about half the mean magnitude of the Ca^2+^ shift observed for CTRL myocytes (**Fig. 7B**. CTRL 2 Hz: 0.664±0.052, CTRL 4 Hz: 0.775±0.083, HFSD 2 Hz: 0.688±0.099, HFSD 4 Hz: 0.741±0.126, F_340:380_, 2-way ANOVA significant interaction effect, **Fig. 7C**. CTRL 0.110±0.051; HFSD 0.054±0.049, F_340:380_). No treatment or interaction effects were observed for systolic Ca^2+^, Ca^2+^ amplitude, or time constant of Ca^2+^ transient decay (**Table. 3**). Frequency elevation was associated with a reduction in diastolic sarcomere length in both CTRL (3%) and HFSD (5%) loaded cardiomyocytes (**Fig. 7D**. CTRL 2 Hz: 1.69±0.04, CTRL 4 Hz:1.64±0.05, HFSD 2 Hz: 1.69±0.04, HFSD 4 Hz: 1.61±0.07 µm). The combination of normal diastolic sarcomere length reduction, concomitant with suppressed diastolic Ca^2+^ increase in HFSD cardiomyocytes, may indicate elevated diastolic responsiveness to Ca^2+^ in the HFSD cardiomyocyte.

**Fig. 7.**
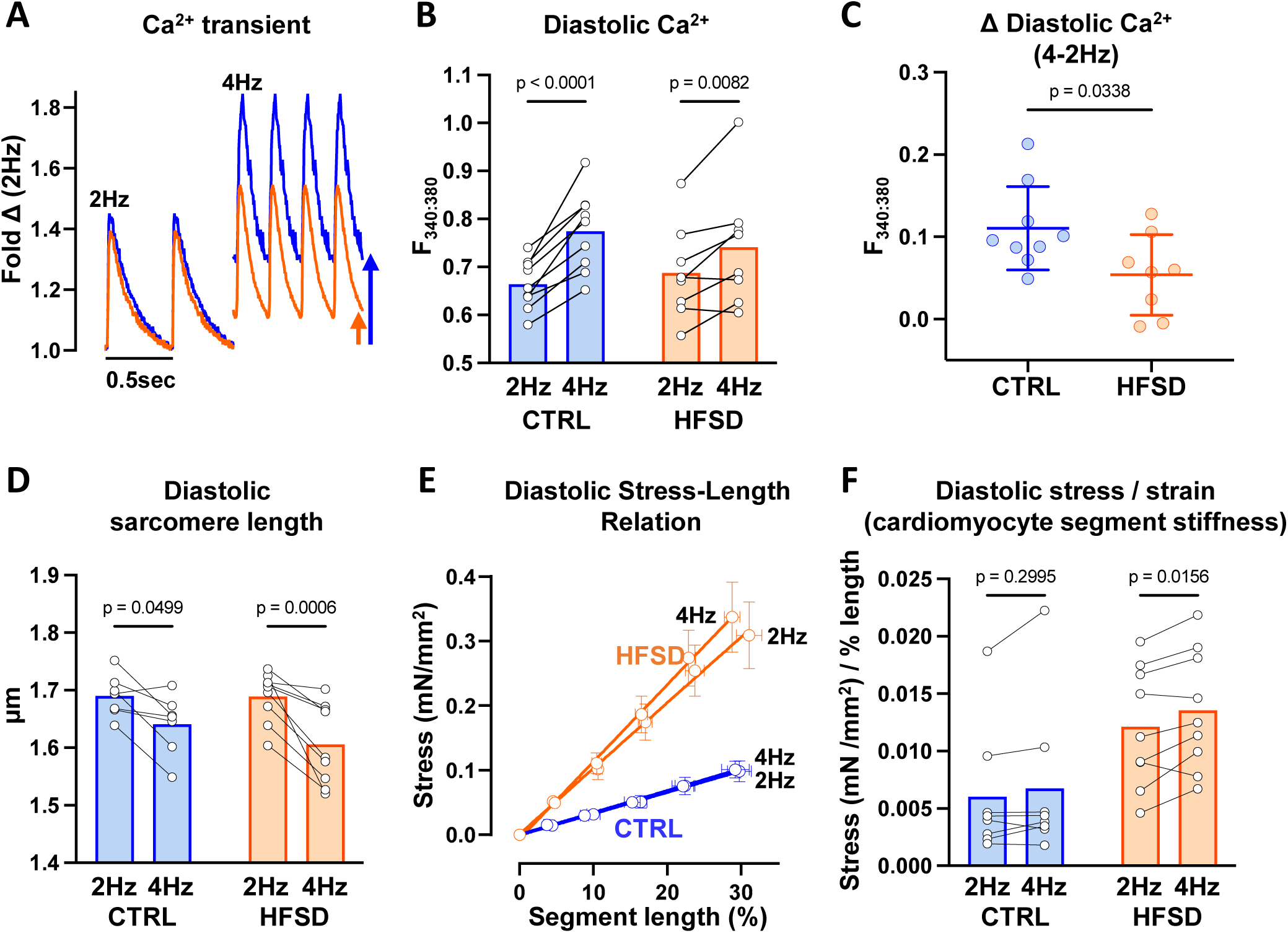
Pacing induced changes in cardiomyocyte cytosolic Ca^2+^ levels and stiffness in CTRL & HFSD. **A.** Exemplar CTRL and HFSD cardiomyocyte Ca^2+^ transients at 2 and 4 Hz pacing. **B.** Diastolic cardiomyocyte Ca^2+^ levels measured before and after transition from 2 to 4 Hz pacing in HFSD & CTRL. **C.** Diastolic cardiomyocyte Ca^2+^ level mean shift with transition from 2 to 4 Hz pacing in HFSD & CTRL. **D.** Diastolic sarcomere length measured before and after transition from 2 to 4 Hz pacing in HFSD & CTRL. **E.** Diastolic mean cardiomyocyte segment stress-length relation (linear regression fit) for HFSD and CTRL at 2 and 4 Hz pacing. **F.** Diastolic mean cardiomyocyte segment stress-length relation slope (mN/mm^2^) normalized relative to segment stretch (% length) yields cardiomyocyte mean segment stress/strain = stiffness [(mN/mm^2^) /strain (% length)] for HFSD and CTRL at 2 and 4 Hz pacing. Comparison between two groups performed using unpaired t-test. With two independent variables a repeated measures 2-way ANOVA Sidaks post-hoc analysis was performed. Data are presented as mean ± standard deviation (C), mean ± standard error of the mean (E) or mean with paired individual points (B, D, F). For **B. & C.** (CTRL: N = 4 mice, n = 9 cells; HFSD: N = 4 mice, n = 8 cells). For **D. & E.** (CTRL: N = 4 mice, n = 7 cells; HFSD: N = 4 mice, n = 9 cells). For **F.** (CTRL: N = 4 mice, n = 8 cells; HFSD: N = 4 mice, n = 9 cells).

**Table 3.**
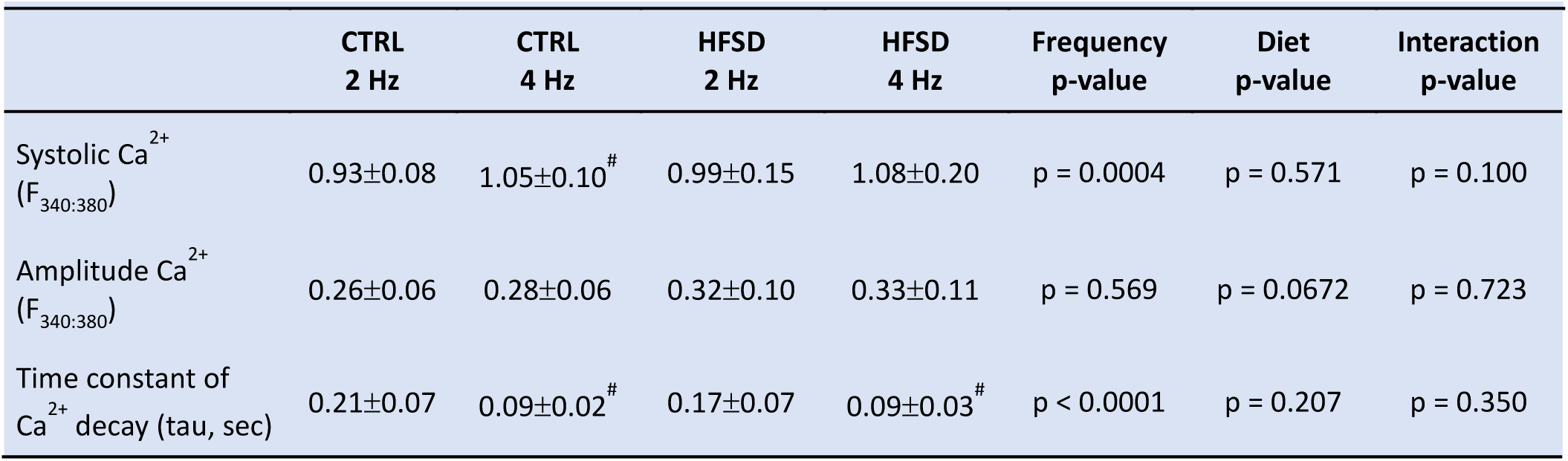
Ca^2+^ transient parameters for loaded CTRL and HFSD mouse cardiomyocytes. Comparisons in data with two independent variable were performed using a mixed analysis 2-way ANOVA with Sidak’s post-hoc analysis (CTRL, 2 Hz: N = 5 mice, n = 13 cells; CTRL, 4 Hz: N = 4 mice, n = 9 cells; HFSD, 2 Hz: N = 5 mice, n = 14 cells; HFSD, 4 Hz: N = 4 mice, n = 8 cells). Data are presented as mean ± standard deviation. P-values for ANOVA effects (Frequency, Diet, Interaction) are listed within the table. #<0.05: Significant pairwise post-hoc frequency differences vs CTRL 2 Hz or HFSD 2 Hz. No pairwise post-hoc treatment differences were observed.

While frequency elevation did not change the stress/strain relation in CTRL cardiomyocytes, a significant increase in the stress/strain gradient was observed with increased frequency in HFSD cardiomyocytes (**Fig. 7E&F**. CTRL 2 Hz: 0.0060±0.0056, CTRL 4 Hz:0.0068±0.0068; HFSD 2 Hz: 0.0121±0.052, HFSD 4 Hz: 0.0136±0.0052 (mN/mm^2^) / % length). This may indicate a larger population of cross-bridges contributing to diastolic stiffness at heightened workloads in HFSD cardiomyocytes although other potential non-cross-bridge sources cannot be excluded. Since Ca^2+^ shift in response to increased pacing is modest (relative to CTRL), increase in stiffness cannot be attributed to rise in Ca^2+^ levels *per se.* Shift in the myofilament-Ca^2+^ response during diastole is a potential alternate explanation.

## DISCUSSION

Here we provide the first quantitative analysis of intact cardiomyocyte stiffness and performance deficits associated with acquired cardiometabolic disease. We demonstrate that a component of cardiac diastolic dysfunction in cardiometabolic disease derives from intrinsic cardiomyocyte mechanical abnormality.

The concept of myocardial stiffness is a frequently invoked qualitative descriptor of the pathophysiologic phenotype characterizing diastolic dysfunction. Clinically and experimentally the echocardiography parameter E/e’ is used as an index of myocardial stiffness in disease diagnosis. E/e’ is a compound measure of cellular and extracellular matrix stiffness contributors. Despite consistent *in vivo* findings, an understanding of the cardiomyocyte-specific contribution to tissue stiffness has been difficult to identify. Using a series of intact single cell recording maneuvers, we have established that the stiffness characteristics of individual isolated cardiomyocytes from normal and dysfunctional hearts can be defined, normalized and compared. Further, a significant relationship was identified between the mean stiffness of single cardiomyocytes and the extent of the origin heart diastolic dysfunction (i.e. E/e’ vs Young’s Modulus [(mN/mm2) / %length]).

To recapitulate, cardiometabolic disease induced by dietary intervention in mice was characterized by marked body weight elevation (50%, ‘obesity’) and glucose intolerance. *In vivo* echocardiography diagnosed significant diastolic dysfunction with a 35% elevation of the E/e’ in HFSD animals in absence of systolic functional deficit. In paced non-loaded conditions, the sarcomere contraction cycles & Ca^2+^ transients (fura-2) were similar in CTRL and HFSD cardiomyocytes. When the same cardiomyocytes were loaded, the shortening extent and maximal lengthening rate were reduced for both diet conditions. In HFSD cardiomyocytes the reduction in deformation rate during lengthening (but not shortening) was accentuated, indicating a selective negative impact of loading on relaxation of the HFSD-derived cardiomyocytes. In HFSD cardiomyocytes there was a larger increase in diastolic force per cross-sectional area (stress: mN/um^2^) in response to standardized step increases in length (strain: % initial length [µm]). From these data a ‘stress/strain’ parameter was determined, defining diastolic stiffness for each myocyte i.e. ‘Youngs Modulus’ ([mN/µm^2^]/[% initial length[µm]). Mean stiffness was 70% higher in cardiomyocytes of HFSD animals. Thus the increase in cardiomyocyte stiffness index was approximately double the elevation of the *in vivo* E/e’ dysfunction index. Using matched *in vivo* and *in vitro* data, a significant correlation was found between diastolic dysfunction (E/e’) and mean cardiomyocyte stiffness value for HFSD-diet animals, while there was no relationship between these parameters in the myocytes of CTRL-diet fed group. To evaluate possible influence of mechanical state on the Ca^2+^ transient, the mean transient decline parameters (tau) from non-loaded, loaded and stretched cardiomyocytes were compared. In HFSD cardiomyocytes the time constant of Ca^2+^ transient decay was increased in stretched cardiomyocytes relative to non-loaded, a feature not observed in CTRL cardiomyocytes. Finally, in response to an *in vitro* pacing increase HFSD cardiomyocytes exhibited accentuated diastolic stiffness even with a smaller increase in diastolic Ca^2+^ level. Collectively, these findings suggest a previously undescribed alteration in Ca^2+^-myofilament interactions indicative of a cross-bridge component to cardiomyocyte stiffness in cardiometabolic disease.

### Dietary model of acquired metabolic syndrome and cardiac diastolic disease

High fat/sugar diet feeding (i.e. ‘western diet’) has previously been shown experimentally to induce diastolic dysfunction in rodents. Elevated E/e’ is a remarkably consistent echocardiographic feature of high fat/sugar diet fed mouse myocardial performance while E/A and IVRT may be elevated, reduced, or unchanged relative to control counterparts (Pulinilkunnil *et al*., 2014; Dia *et al*., 2020; Sowers *et al*., 2020; Wingard *et al*., 2021; Daniels *et al*., 2022). The dietary regime employed in this study has been well characterized as a model of diastolic dysfunction, with gradual development of metabolic disturbance (stable rate of weight gain and mild hyperglycemia), with maintained systolic function (ejection fraction and fractional shortening). The data presented here are consistent with clinical observations linking cardiometabolic disease and diastolic dysfunction.

As we have reported previously, over the time span of the dietary intervention differential fibrosis is not observed (Daniels *et al*., 2022; Wells *et al*., 2023). The finding of increased stiffness in cardiometabolic disease is consistent with some aspects of prior experiments involving animal feeding of obesogenic diets, using other rodent preparation types – including permeabilized cardiomyocytes, tissue strips/papillary muscles, and non-paced cardiomyocytes (Leopoldo *et al*., 2010; Gonçalves *et al*., 2016; Sowers *et al*., 2020). Interestingly stiffness in those studies was attributed to titin or extracellular matrix properties. Our protocol with intact single cardiomyocytes allows a mechanistic extension of these previous findings.

### Loading the intact cardiomyocyte - functional deficits revealed in HFSD

Non-loaded contracting cardiomyocytes of CTRL and HFSD mice exhibited identical shortening-lengthening cycles, but a distinctly different decrement in lengthening rate when exposed to load (i.e. auxotonic with progressive ‘pre-load’ increase). Our findings show unequivocally that cardiomyocyte mechanical dysfunction cannot be detected by analysis of non-loaded shortening. With loading, the shortening extent and the maximal lengthening rates were reduced for both CTRL and HFSD. In HFSD the lengthening rate reduction was accentuated even when shortening extent was accounted for. These findings indicated specific influence of loading on lengthening in HFSD cardiomyocytes and could reflect impaired subcellular relaxation processes specific to the HFSD myocardium.

Others have demonstrated a physiological effect of loading on shortening in cardiac cells/tissues in different species (using slow pacing and sub physiologic temperature conditions), although the pronounced effect on relaxation is not consistently reported (Lab *et al*., 1984; White *et al*., 1995; Shimkunas *et al*., 2021). This is consistent with *in vivo* human clinical observations where there is a strong inverse association between late-systolic loading and left ventricular wall displacement velocity during early diastole (Borlaug *et al*., 2007). Importantly, increased dependence of relaxation on after-load is reported to be an *in vivo* feature of heart failure with diastolic dysfunction (Eichhorn *et al*., 1992; Gillebert *et al*., 2000). Regarding the Ca^2+^ transient, in the current study a subtle prolongation effect is observed in both CTRL and HFSD cardiomyocytes when loaded (but not stretched) which is consistent with recently reported findings in rat LV trabeculae (Dowrick *et al*., 2022). The variability in the sarcomere shortening recorded for both non-loaded and loaded cardiomyocytes is notable but consistent with other studies assessing relaxation of rodent cardiomyocytes (Delbridge & Roos, 1997; Clark & Campbell, 2019). A range of non-loaded cardiomyocyte mean diastolic sarcomere lengths are reported in the literature. When cells are evaluated *in vitro* and not constrained by local structural elements sarcomere lengths <1.8µm are commonly observed and consistent with our findings (Lim *et al*., 2000; Bub *et al*., 2010). *In vitro* experimental conditions can also contribute to variation in diastolic sarcomere lengths (e.g. pacing frequency, temperature and buffer tonicity) (Roos *et al*., 1982; Chung & Campbell, 2013; Khokhlova *et al*., 2022).

### Characterizing cardiomyocyte stretch & pacing challenge responses

In order to quantify intact single cardiomyocyte stiffness our approach was to apply a calibrated stretch protocol, based on pre-measurement of a stretch calibration constant for each myocyte, tracking changes in force generation and cytosolic Ca^2+^ responses for each stretch step. This calibrated step-by-step approach enabled the stress and strain data to be gathered from actively paced cardiomyocytes over extended periods (i.e. 80-100 sec) and computed to generate a definitive, comparative stiffness parameter for each cardiomyocyte.

In the intact myocyte model, the internal cytosolic regulatory milieu is preserved, and the cytoskeletal/sarcomeric/organelle structural framework is integral. Thus contributions to diastolic ‘stiffness’ reflect components of the structural passive resistance to deformation and the basal crossbridge-derived activity. Recent studies of permeabilized cardiac muscle strip bundles have explored the separate contributions of mictrotubule, titin, actin-associated, sarcolemmal and desmin-linked components to passive force development (Loescher *et al*., 2023). Prior studies of ‘skinned’ or quiescent myocardial preparations have also identified sources of passive stiffness elevation, with titin isoform switch and modification identified as major modulators of stiffness in cardiometabolic disease (Leopoldo *et al*., 2010; Hamdani *et al*., 2013; Gonçalves *et al*., 2016; Hopf *et al*., 2018; Sowers *et al*., 2020; Abdellatif *et al*., 2021).

Our findings indicate involvement of a cytosolic Ca^2+^ component of the stretch response. Ca^2+^ fluorescence analyses from these stretch protocols showed novel stretch-dependent prolongation of Ca^2+^ transient decay in the HFSD cardiomyocytes. The time constant of Ca^2+^ transient decay increased with loading and further with stretching – a feature not evident in CTRL cardiomyocytes and which provides additional mechanistic insight.

The final pacing frequency protocol was implemented as a relative workload stress to the loaded cardiomyocyte. With a doubling of pacing frequency, a significant increase in the stress/strain gradient was observed in HFSD cardiomyocytes (and not CTRL), and a similar reduction in diastolic sarcomere length in both CTRL and HFSD was observed. While cardiomyocyte diastolic Ca^2+^ level increased in both CTRL and HFSD – the HFSD increase was less marked than CTRL. This was a surprising finding, not previously reported in any other model setting. A suggestion of Ca^2+^ hyper-sensitivity arises (Davis *et al*., 2007) amongst other explanations to consider.

### Mechanistic insights into cross-bridge derived stiffness

Here we report that a cardiometabolic disease phenotype induced by obesogenic diet includes a 70% increase in cardiomyocyte stiffness which contributes to a 35% increase in diastolic dysfunction *in vivo* (by echo E/e’). Several lines of evidence emerging from this study suggest that Ca^2+^-related mechanism/s may contribute to cardiomyocyte stiffness increase. Specifically, in the loaded HFSD cardiomyocyte (vs CTRL):

a. load confers accentuated lengthening rate deficit
b. with stretch, the Ca^2+^ transient decay time course is prolonged
c. with more rapid pacing, relatively small increases in diastolic Ca^2+^ are associated with elevated stiffness Together these cardiomyocyte phenotypes would be consistent with selective impairment of relaxation rate, selective myofilament hyper-activation, impeded diastolic filling and reduced diastolic reserve.

This combination of observations is consistent with the notion that stretch-dependent cross-bridge attachment is prolonged in HFSD cardiomyocytes (or conversely detachment is delayed). We can speculate that an alteration in the 3-dimensional geometry of the low affinity Ca^2+^ binding pocket in troponin C (TnC) would produce this effect, by stabilizing Ca^2+^ in the pocket for longer every activation cycle. This could increase TnC buffering power which would explain the relatively small Ca^2+^ rise in HFSD in response to increased pacing (Eisner *et al*., 2023). A manifestation of this altered binding function could be apparent Ca^2+^ hypersensitivity. The TnC-Ca^2+^ dissociation rate has been theorized as an effector of the transient decay time-course (Robinson *et al*., 2018). A role for the troponin complex in regulation of Ca^2+^ affinity is well established (Chemla *et al*., 2000). Given the load and stretch-dependent nature of these findings, a role for stress sensor proteins such as titin is likely. Indeed, there is some evidence to suggest that titin isoform regulates TnC low-affinity binding pocket opening and Ca^2+^ affinity in response to stretch via regulating the number of attached cross-bridges (Li *et al*., 2019). Further evidence of structural modifications of specific myofilament regions which form and regulate this pocket is required. Mass spectrometry studies by us and others provide proof of concept of relevant post-translational modification types (Janssens *et al*., 2018; Loescher *et al*., 2022).

### Limitations

Given the diastolic dysfunction focus of this study, investigations have primarily examined cardiomyocyte relaxation and lengthening behaviors. Diastolic and systolic interactions are important and additional protocols are in development for these studies. In this regard, we defer to the extensive body of technologic research undertaken to develop software and instrumentation to perform single cardiomyocyte ‘Pressure-Volume (PV) loops’ (Helmes *et al*., 2016). Using this approach, to mimic PV cycles in the intact myocardium, stretch is applied after relaxation has occurred by using a force clamp to maintain a predefined diastolic force level. This approach has been used to discover that titin truncation shifts the working sarcomere length range preserving diastolic stiffness at the expense of systolic performance (Methawasin *et al*., 2022) and that highly compliant titin allows cardiomyocytes to function at high diastolic Ca^2+^ levels (Najafi *et al*., 2019). Potentially, this approach could also be used to evaluate the temporal relationship between slow or incomplete relaxation and cardiomyocyte cross-bridge derived stiffness in a semi-situ manner. Operator expertise is imperative with this system which to date has been very productive in the hands of the developers.

In our work we have adopted a pragmatic approach, simpler to implement – although still very challenging. Our protocol involves the application of cumulative ramp steps with about 20% non-loaded-to-loaded myocyte conversion success rate. When animals have many months of dietary treatment history, and as cell isolation from older and diseased hearts is difficult, an optimized loading and stretching approach is key. The two approaches have not been benchmarked, although it can be predicted that the viscous components of stiffness are likely underestimated when stretches are slow and relatively small such as in the current study (Caporizzo & Prosser, 2021). The capacity to directly and reproducibly determine Young’s Modulus (stress/strain) with a slow stretch routine has useful comparative value. An additional aspect of our pragmatic approach has been to evaluate cardiomyocyte segment stretches as average sarcomere stretch methods can be challenging to track optically.

With ongoing research, a priority is to undertake longitudinal studies, including both sexes, to investigate the relative timing of the emergence of *in vivo* and *in vitro* stiffness phenotypes as differences in Ca^2+^ handling have previously been observed between control male and female rodents (Farrell *et al*., 2010). This has high translational relevance as we have previously identified a differential time course of diastolic dysfunction emergence in females and males, with increased female vulnerability (Reichelt *et al*., 2013; Chandramouli *et al*., 2018).

### In conclusion

This is the first study to link elevated intact single cardiomyocyte stiffness with *in vivo* diastolic dysfunction in a dietary acquired cardiometabolic disease mouse model. The novel findings identify augmented load dependent TnC-Ca^2+^ interactions as one potential feature of myofilament function which contributes to cardiomyocyte stiffness in hearts exhibiting cardiometabolic diastolic dysfunction. Taken together, the findings emphasize the importance of mechanical loading and pacing frequency in revealing latent diastolic abnormalities at the cardiomyocyte level. The rationale for further studies identifying specific subcellular and molecular mechanisms of cardiomyocyte stiffness in cardiometabolic disease prior to failure onset is compelling.

## BIOGRAPHY

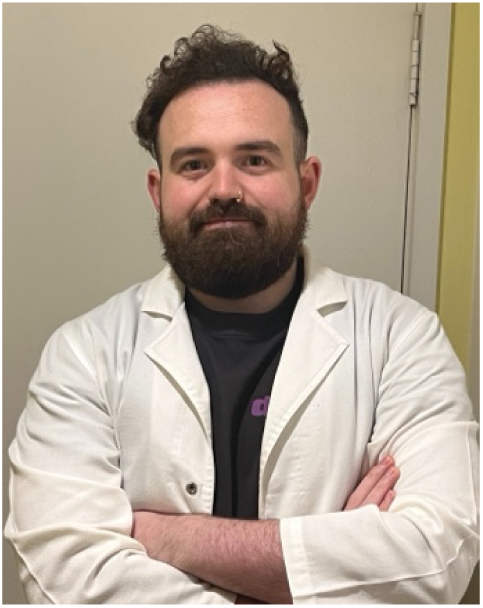

**Johannes Janssens** completed his PhD in the Department of Anatomy and Physiology at the University of Melbourne (Australia) where the current work was performed. After completing his PhD he received a Fulbright Postdoctoral Scholarship to develop a novel blood biomarker of diabetic heart disease at Cedars-Sinai Medical Center (Los Angeles, CA) where he continues to work currently. Johannes’ research aspirations involve linking large-scale proteomic features of hearts and cardiomyocytes with their functional phenotype to develop novel therapeutic and diagnostic strategies for cardiac pathophysiology. In particular diseases where diastolic dysfunction is a prominent characteristic.

## ADDITIONAL INFORMATION

### Data availability statement

The data that support the findings of this study are available from the corresponding author upon reasonable request.

### Author contributions

All experiments were performed at the Cardiac Phenomics Laboratory at the University of Melbourne, Australia. J.V.J., K.M.M. and L.M.D.D. contributed to the conception and research design of the work. All authors contributed to the acquisition, analysis or interpretation of data for the work and contributed to drafting the work or revising it critically for important intellectual content. All authors provide final approval of the version to be published and agreement to be accountable for all aspects of the work.

